# Optimizing targeting of pinyon-juniper management for sagebrush birds of conservation concern while avoiding imperiled pinyon jay

**DOI:** 10.1101/2022.07.21.500993

**Authors:** Jason R. Reinhardt, Jason D. Tack, Jeremy D. Maestas, David E. Naugle, Michael J. Falkowski, Kevin E. Doherty

## Abstract

Contemporary restoration and management of sagebrush-dominated (*Artemisia* spp.) ecosystems across the intermountain west of the United States increasingly involves the removal of expanding conifer, particularly juniper (*Juniperus* spp.) and pinyon pine (*Pinus edulis, P. monophylla*). The impetus behind much of this management has been the demonstrated population benefits of sagebrush restoration via conifer removal to greater sage-grouse (*Centrocercus urophasianus*), a species of conservation concern. One of the challenges with scaling up from a focal-species approach to a community-level perspective, however, is balancing the habitat requirements of different species, some of which may overlap with sage-grouse and others which may have competing habitat needs. Here, we use a systematic conservation planning approach to compute spatial optimizations which prioritize areas for conifer removal across the sage-grouse range while incorporating woodland and sagebrush songbirds into decision-making. Three of the songbirds considered here, Brewer’s sparrow (*Spizella breweri*), green-tailed towhee (*Pipilo chlorurus*), and sage thrasher (*Poocetes gramineus*), are sagebrush-obligates, while another is a woodland-obligate, the pinyon jay (*Gymnorhinus cyanocephalus*). We find that the inclusion of sagebrush-obligates expands the model-selected area of consideration for conifer management, likely because habitat overlap between sagebrush-obligates is imperfect. The inclusion of pinyon jay, a woodland-obligate, resulted in substantial shifts in the distribution of model-selected priority areas for conifer removal – particularly away from pinyon jay strongholds in Nevada and east-central California. Finally, we compared the conifer optimizations created here with estimates of ongoing conifer removal efforts across the intermountain west and find that a small proportion (13-18%) of management efforts had occurred on areas predicted as being important for pinyon jay, suggesting that much of the ongoing work is already successfully avoiding critical pinyon jay habitat areas.

## Introduction

Conifer expansion into sagebrush-dominated (*Artemisia* spp.) ecosystems is an ongoing phenomenon that has garnered attention in recent years (Miller et al. 2008; Miller et al. 2011; Chambers et al. 2017; Miller et al. 2019). The extent of native juniper (*Juniperus* spp.) and pinyon pine (*Pinus edulis, P. monophylla*) woodlands in western North America has increased substantially since 1860, with cover increases ranging between 125% and 625% depending on the region (Miller et al. 2008). Much like other instances of woodland expansion into grass- and shrub-dominated systems from around the world (see Nackley et al. 2017), pinyon-juniper expansion into sagebrush systems is a concern due to the associated changes in plant diversity (Miller et al. 2000; Bates 2005), ecosystem resistance and resilience (Chambers et al. 2007; Chambers et al. 2014; Miller et al. 2014), and impacts on native wildlife (Rickart et al. 2008; Baruch-Mordo et al. 2013; Severson et al. 2016). The ongoing impact of pinyon-juniper expansion on these different aspects of sagebrush ecosystems has elicited a strong interest in restoration and management efforts focused on reducing and removing expanding pinyon-juniper woodlands across the U.S. Intermountain West (Miller et al. 2017, Maestas et al. 2021).

Much of the pinyon-juniper management being conducted in sagebrush ecosystems has been driven by conservation interest in the greater sage-grouse (*Centrocercus urophasianus*, hereafter ‘sage-grouse’), largely because this imperiled species was a candidate for listing under the U.S. Endangered Species Act (USFWS 2015). Recent management activities have largely focused on removing pinyon-juniper from sites with relatively low tree canopy cover, in response to research conducted by Baruch-Mordo et al. (2013), which found the sage-grouse to be sensitive to even low levels of conifer cover (∼3-4%) on breeding sites. This revelation has spurred both management activities and motivated numerous studies assessing pinyon-juniper removal as an integral component of sage-grouse habitat improvement (e.g., Prochazka et al. 2017; Severson et al. 2017a,b; Olsen et al. 2021; Rabon et al. 2021). Much of this recent work reinforces the efficacy of pinyon-juniper removal as a tool for sage-grouse conservation via a number of different metrics, including increasing nesting habitat availability, space-use, bird abundance, and population growth (Severson et al. 2017a; Sandford et al. 2017; Severson et al. 2017b, Olsen et al. 2021).

Several broad-scale spatial datasets and decision-support tools have been developed in tandem with the growth in sage-grouse and pinyon-juniper management activities and research. One of the most fundamental of these is a high-resolution (1-m^2^) conifer cover map (Falkowski et al. 2017) which provides a synoptic view of pinyon-juniper across the 458,000 km^2^ sage-grouse range. This dataset has been used by land managers and stakeholders as a planning tool, and in conjunction with other data, has informed a number of research efforts, including assessments of the impact of conifer expansion on sage-grouse nesting site selection (Severson et al. 2017a) and developing frameworks for considering local-scale conifer cuts in a landscape context (Reinhardt et al. 2017). Broad scale datasets and analyses such as this have also proven essential for evaluating ongoing conifer management efforts (Reinhardt et al. 2020). Recently developed tools such as the Rangeland Analysis Platform (RAP; Jones et al. 2018, Allred et al. 2021) expand the spatial toolset even further and make data available to managers and scientists by incorporating a wide range of vegetative cover types across large timescales.

The expansion of these large-scale spatial analyses and tools to include other species and community components is encouraging because while the sage-grouse has been much of the impetus behind conifer removal efforts in sagebrush systems, several other species have been shown to benefit from conifer management (Maestas et al. 2021). Removal of conifers from sagebrush systems has been shown to result in more herbaceous plant cover and biomass (Bates et al. 2017, Fick et al. 2022), increases in Brewer’s sparrow (*Spizella breweri*), green-tailed towhee (*Pipilo chlorurus*), and vesper sparrow (*Poocetes gramineus*) populations (Crow and Van Riper 2010; Donnelly et al. 2017; Holmes et al. 2017), enhanced mule deer (*Odocoileus hemionus*) fawn overwintering survival (Bergman et al. 2014a), and possibly mule deer size and condition (Bender et al. 2013; Bergman et al. 2014b).

While sage-grouse – acting as a focal-species driver of sagebrush ecosystem restoration has been beneficial for a number of species, it is also important to consider the possible non-target negative impacts of what has effectively been an umbrella species approach to management (Rowland et al. 2006, Zeller et al. 2021). Indeed, there has been an ongoing debate regarding whether pinyon-juniper removal efforts in support of sage-steppe improvement projects is resulting in substantial loss of pinyon-juniper woodlands, and negatively impacting woodland-associated species (Boone et al. 2018; Coe et al. 2018; Maestas et al. 2019; Clark et al. 2019, Zeller et al. 2021). A recent review (Bombaci and Pejchar 2016) found little overall negative impact of conifer removal on woodland associate species, but data were lacking for some taxa, particularly invertebrates. One species of particular concern with regard to conifer removal is the pinyon jay (*Gymnorhinus cyanocephalus*). Pinyon jay populations have demonstrated significant long-term population declines (Sauer et al. 2017) and is a species of high conservation concern (Boone et al. 2018). Given pinyon jay reliance on pinyon pine woodlands, there has been some concern that sagebrush management efforts incorporating conifer removal may adversely affect the species. Despite these concerns, however, there has been relatively little work has been done to examine potential mechanisms behind the pinyon jay’s population decline or effects of woodland management relative to drought or other ecosystem stressors (Boone et al. 2018).

Given the potential multi-species implications of conifer removal, it is important for management planning tools and datasets to incorporate species with potentially overlapping and competing habitat requirements – beyond sage-grouse alone. Here, we use a systematic conservation planning approach implemented as a spatial optimization scenario to target conifer removal in a way that balances the habitat requirements of multiple bird species. Specifically, we set out to optimize areas for conifer removal based on the habitat requirements of the greater sage-grouse, three sagebrush-obligate songbirds: the Brewer’s sparrow, sage thrasher (*Oreoscoptes montanus*), and green-tailed towhee, and the woodland-obligate pinyon jay. Each of the sagebrush-obligate songbirds and the pinyon jay were selected because they demonstrate long-term population declines based on North America Breeding Bird Survey (BBS) data (Sauer et al. 2017; Tack et al. *in review*).

Our objectives in this work were to: 1) assess the relative effect of incorporating different types of species habitat requirements on optimal locations for conifer removal across the sage-grouse range, 2) evaluate in detail the effect of incorporating pinyon jay – a woodland obligate with potentially competing habitat requirements – on conifer removal optimization, and 3) determine how recent conifer removal efforts align with the different optimizations, particularly as it relates to areas potentially important to pinyon jay.

## Methods

### Study Area

We considered the same region of the intermountain west for which canopy cover was mapped by Falkowski et al. (2017), which includes parts of Oregon, Idaho, Montana, California, Nevada, Arizona, Wyoming, and Colorado. This region overlaps with Western Association of Fish and Wildlife Agencies Sage-Grouse Management Zones (SGMZ) III, IV, V, and VII, covering approximately 458,000 km^2^, 134,000 km^2^ of which has at least some conifer cover (i.e., > 0%). A substantial portion (250,971 km^2^) of the study area falls within U.S. Fish and Wildlife Service Priority Areas for Conservation (PACs) for sage-grouse (U.S. Fish and Wildlife Service, 2013).

The study area is dominated by sagebrush (*Artemisia* spp.) ecosystems, Pinyon pine and juniper woodlands, and agricultural land (Miller et al. 2008; Homer et al. 2015). The area of interest encompasses a large portion of the intermountain west where climate conditions tend to be cold and dry (Kottek et al. 2006). Summer (June-August) temperatures generally range from a maximum of 11.3 to 37.3 °C to minimums of -1.2 to 17.1 °C, while winter (December-February) temperatures range from a maximum of -6.2 to 11.5 °C with a minimum of -19.5 to -1.8 °C (Daly et al. 2008). Summer precipitation averages between 16 and 212 mm, while winter averages between 13 and 532 mm (Daly et al. 2008).

### Data

A series of geospatial datasets were required to conduct the optimization analyses. To identify where pinyon-juniper woodlands were located on the landscape in our study area we leveraged the high-resolution conifer cover prediction map created by Falkowski et al. (2017). This dataset maps conifer at a 1-m spatial resolution and was created using a spatial wavelet analysis approach (Falkowski et al. 2006) on National Agricultural Imagery Program (NAIP) aerial imagery. Areas important to sage-grouse were identified using a sage-grouse distribution model created by Doherty et al. (2016) using a random forest (RF) modeling approach in conjunction with numerous environmental considerations.

Habitat suitability models for each of the songbirds included in this analysis (Brewer’s sparrow, green-tailed towhee, sage thrasher, and pinyon jay) were obtained from the U.S. Fish & Wildlife Service (Tack et al. *in review*). Each model was created using a RF modeling approach in conjunction with environmental data and North American Breeding Bird Survey (BBS) data (Sauer et al. 2017).

All spatial data were aggregated by computing zonal statistics across a hexagonal grid with 16.24 km^2^ grid cells (2.5 km on a side). The relatively coarse scale of data aggregation reflects our interest in broad landscape-level patterns rather than fine-scale details.

### Analysis

We used an integer linear programming (ILP) approach to compute the optimization of areas for pinyon-juniper removal across six different sage-grouse and songbird data combinations (Table 1). Brewer’s sparrow was considered in a scenario configuration alongside sage-grouse on its own and in the ‘all sagebrush-obligate songbirds’ group as it is the sagebrush-obligate with the largest population declines among those included in this study (Tack et al. *in review*). The ILP optimization was computed using the *prioritizr* package in R (Hanson et al. 2017) and leveraged the *Gurobi* solver (Gurobi Optimization 2017). Optimizations were run using a minimum set objective which seeks to minimize a ‘cost’ parameter while meeting goals for ‘conservation features’ (Beyer et al. 2016; Hanson et al. 2017). The hexagonal grid over which the spatial data were aggregated was the spatial unit of the optimization, and sage-grouse along with the sagebrush obligate songbirds were the conservation features. Cost was a function of canopy cover for optimization scenarios not including pinyon jay, and canopy cover + pinyon jay habitat suitability for optimizations which included the species – such that grid cells with high canopy cover and high pinyon jay habitat suitability represented high-cost areas. Grid cells that contained no conifer cover were excluded from analyses as they contained no conifer to prioritize for removal. We set proportional targets of 0.3 for each conservation feature, and we used a boundary penalty of 0.001 to discourage heavily fragmented solutions (Ardron et al. 2010).

**Table 1.** Six different sage-grouse and songbird data configurations used to generate conifer removal optimizations.

| Optimization Scenario Configuration | Label |
| --- | --- |
| Sage-grouse only | SG only |
| Sage-grouse, Brewer's sparrow | SG + BRSP |
| Sage-grouse, all sagebrush-obligate songbirds | SG + SBSong |
| Sage-grouse, pinyon jay | SG only + PIJA |
| Sage-grouse, Brewer's sparrow, pinyon jay | SG + BRSP + PIJA |
| Sage-grouse, all sagebrush-obligate songbirds, pinyon jay | SG + SBSong + PIJA |

We generated a portfolio of 100 distinct solutions for each optimization scenario; increasing the portfolio beyond 100 in preliminary runs had no significant impact on overall results or trends. Selection frequency (‘sf’), or the proportion of solutions in which a particular grid cell was selected, was used as an indicator of relative importance (Carwardine et al. 2007; Ardron et al. 2010). Selection frequency was mapped for all optimizations and the distributions were summarized using kernel density plots. Kolmogorov-Smirnov tests were used alongside linear models to evaluate the potential impacts of adding pinyon jay as a consideration to each of the three sagebrush obligate combinations. optimization results were compared between states using density plots, and within states using linear models.

Differences between conifer optimizations which included and excluded pinyon jay were computed using simple zonal subtraction. These differences in priority as a result of the inclusion of pinyon jay were compared with estimates of recent conifer removal efforts attributable to management as predicted by Reinhardt et al. (2020). Areas of conifer removal were sorted into categories based on whether they overlapped areas where the inclusion of pinyon jay caused a loss in selection frequency (i.e., areas that may be particularly important to pinyon jay), a gain in selection frequency (i.e., areas that may be less important to pinyon jay but important to the other species in the optimization), or areas of little to no change (Δsf < ±0.5 standard deviation).

## Results

The conifer optimizations identified between 4,676 and 6,999 grid cells (16.24 km^2^) as being selected in at least one solution (sf > 0) for the entire study area, with the sage-grouse only and sage-grouse + pinyon jay optimizations having the fewest number of selected cells, and all four of the optimizations incorporating additional sagebrush obligates demonstrating over selected 6,000 cells (Table 2).

**Table 2.**
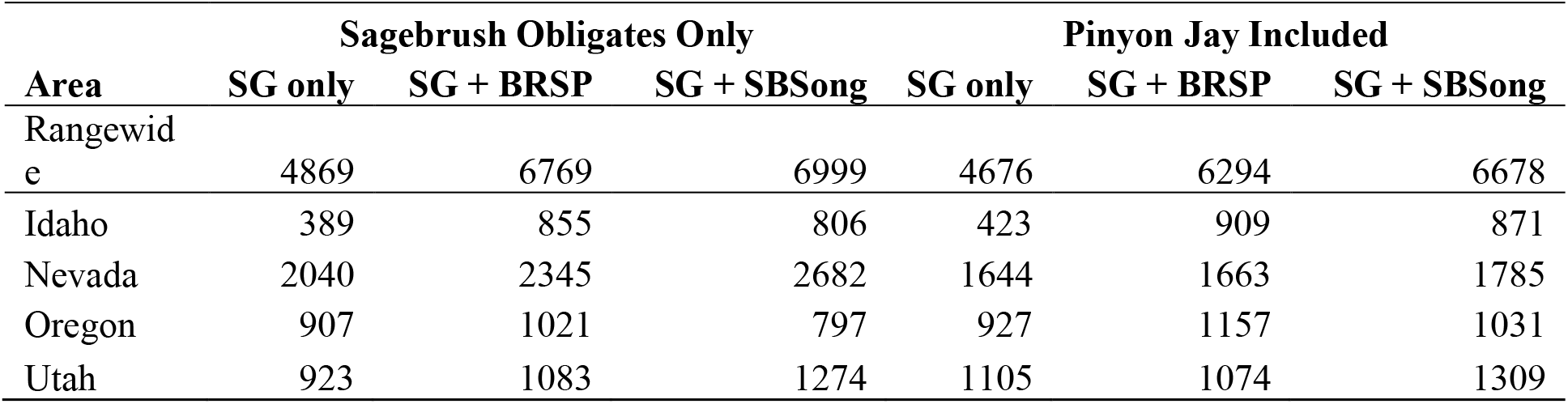
Number of 16.24 km^2^ grid cells selected at least once for each model. Results are summarized rangewide and within four influential states containing the most conifer (Figure 1).

Adding pinyon jay to any optimization scenario resulted in appreciable changes in selection frequency distribution across states (Table 2). Broadly, the inclusion of pinyon jay resulted in a reduction of selected areas in Nevada and an increase in selected areas in Oregon, and to a lesser extent, Idaho and Utah (Table 2). The largest change was found in the sage-grouse + all sagebrush obligate songbird optimization, where adding pinyon jay resulted in a decrease of approximately 900 grid cells in Nevada (Table 2, Figure 1).

**Figure 1.**
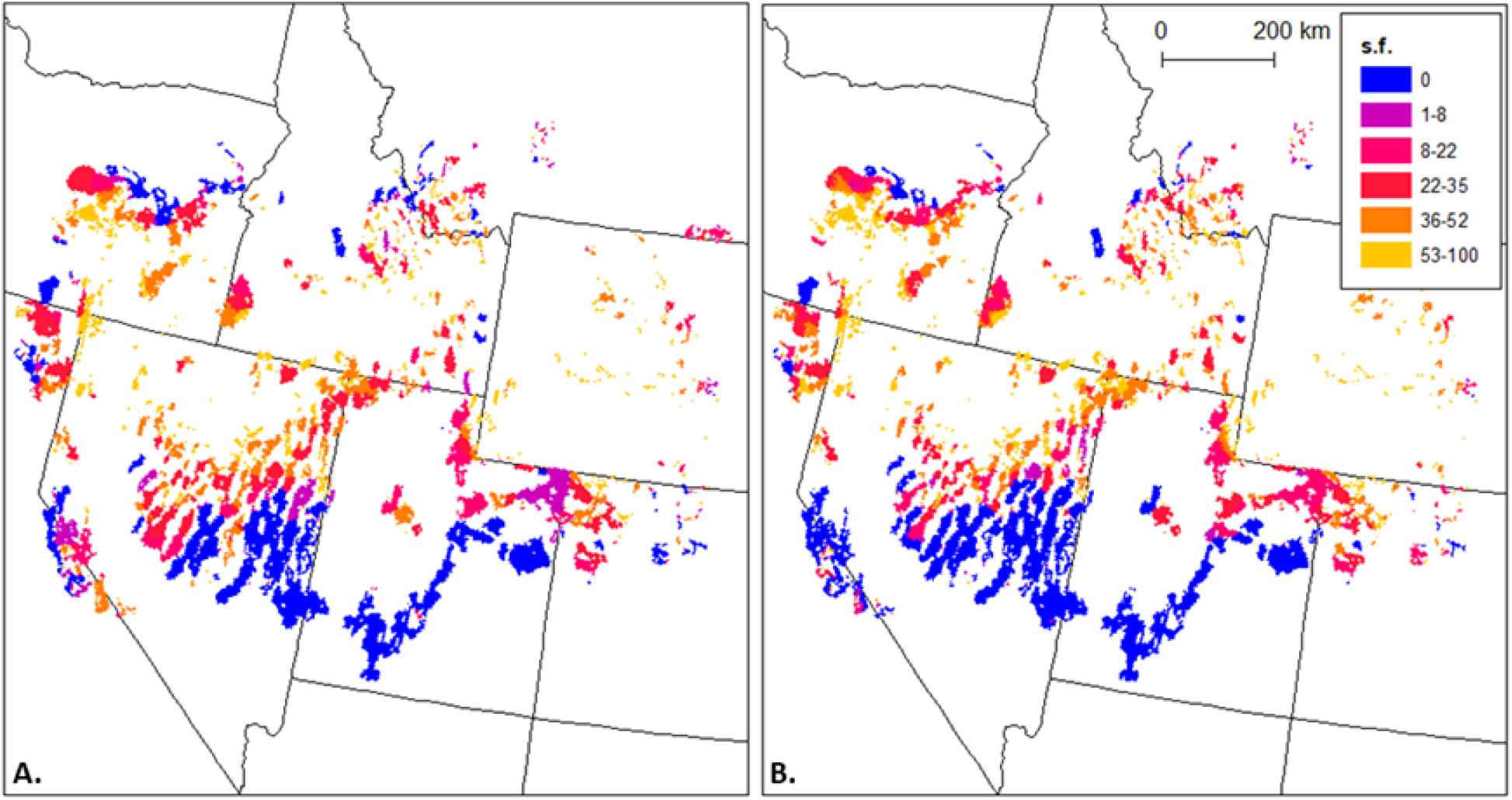
Conifer optimizations for sage-grouse + all sagebrush-obligate songbird combinations. Optimization with (A) and without (B) pinyon jay. Warmer colors depict higher selection frequency (s.f.) indicating the most optimal places for conifer removal. Each map grid cell is a 16.24 km^2^ hexagon, 2.5 km on a side.

Pinyon jay also had an impact on the overall selection frequency distributions, resulting in significant changes in terms of how often particular areas were selected, given that they were selected at all (Figure 2). Indeed, while we found coarse overall relationships between the selection frequencies of models with and without pinyon jay (sage-grouse only *r*^2^ = 0.803; sage-grouse + brewer’s sparrow *r*^2^ = 0.884; sage-grouse + all sagebrush obligates *r*^2^ = 0.783), the inclusion of pinyon jay resulted in meaningful changes to the overall shape of the distribution (sage-grouse only *D* = 0.059, *P* < 0.001; sage-grouse + brewer’s sparrow *D* = 0.077, *P* < 0.001; sage-grouse + all sagebrush obligates *D* = 0.078, *P* < 0.001) (Figure 2).

**Figure 2.**
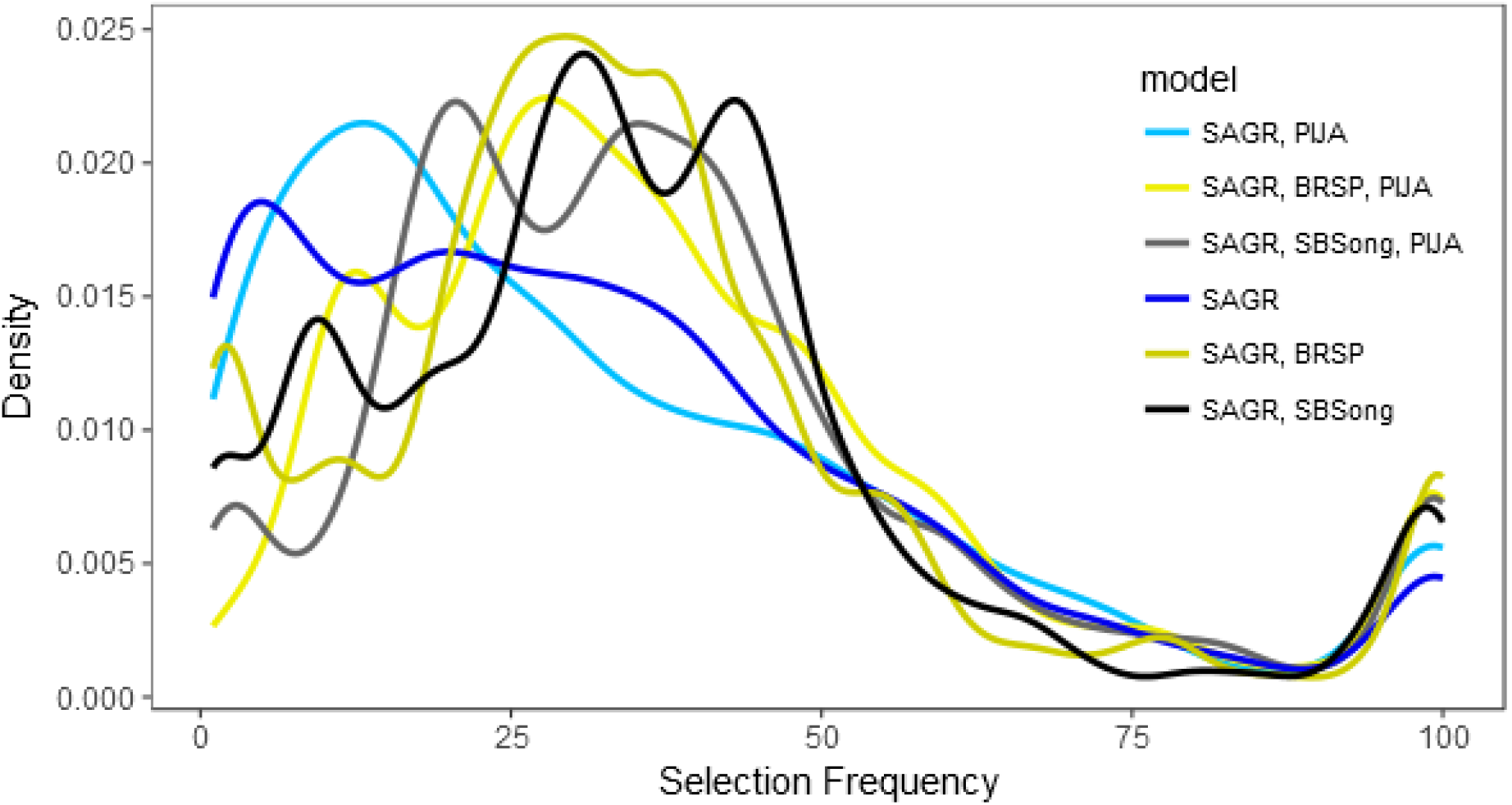
Distributions of selection frequencies for kernel density plots across the study area (Figure 1). Color pairings for six bird groupings depict models with (lighter) and without pinyon jay (darker).

In addition to the quantifiable impact of including pinyon jay in the conifer optimizations at the sage-grouse range wide scale, we also found apparent impacts at the state level, particularly for the four states (Figure 1) with the most prioritized conifer in our analyses. While linear relationships were generally apparent between the selection frequencies of optimizations with and without pinyon jay (ranging from r^2^ = 0.693 to r^2^ = 0.952; Table 3), the inclusion of the pinyon-juniper woodland obligate species had an impact on the overall shape of the selection frequency distributions across each state in all instances except the sage-grouse + all sagebrush obligate optimization for Idaho (*D* = 0.087, *P* = 0.092) (Table 3, Figure 3).

**Table 3.** Impact of adding pinyon jay (PIJA) into optimizations, as measured by differences in distributions of selection frequencies (s.f.) across the range of sage-grouse and within Nevada, Oregon, Utah and Idaho. Differences in s.f. distributions were assessed via the Kolmogorov-Smirnov test.

| Scope | Model set | *D* | *p* | *r*<sup>2</sup> |
| --- | --- | --- | --- | --- |
| Rangewide | SG only | 0.059 | < 0.001 | 0.803 |
|  | SG + BRSP | 0.077 | < 0.001 | 0.884 |
|  | SG + SBSong | 0.078 | < 0.001 | 0.783 |
| NV | SG only | 0.118 | < 0.001 | 0.871 |
|  | SG + BRSP | 0.760 | < 0.001 | 0.768 |
|  | SG + SBSong | 0.096 | < 0.001 | 0.795 |
| OR | SG only | 0.117 | < 0.001 | 0.952 |
|  | SG + BRSP | 0.070 | 0.009 | 0.892 |
|  | SG + SBSong | 0.128 | < 0.001 | 0.871 |
| UT | SG only | 0.167 | < 0.001 | 0.827 |
|  | SG + BRSP | 0.219 | < 0.001 | 0.838 |
|  | SG + SBSong | 0.277 | < 0.001 | 0.800 |
| ID | SG only | 0.087 | 0.092 | 0.770 |
|  | SG + BRSP | 0.099 | < 0.001 | 0.693 |
|  | SG + SBSong | 0.096 | < 0.001 | 0.731 |

**Figure 3.**
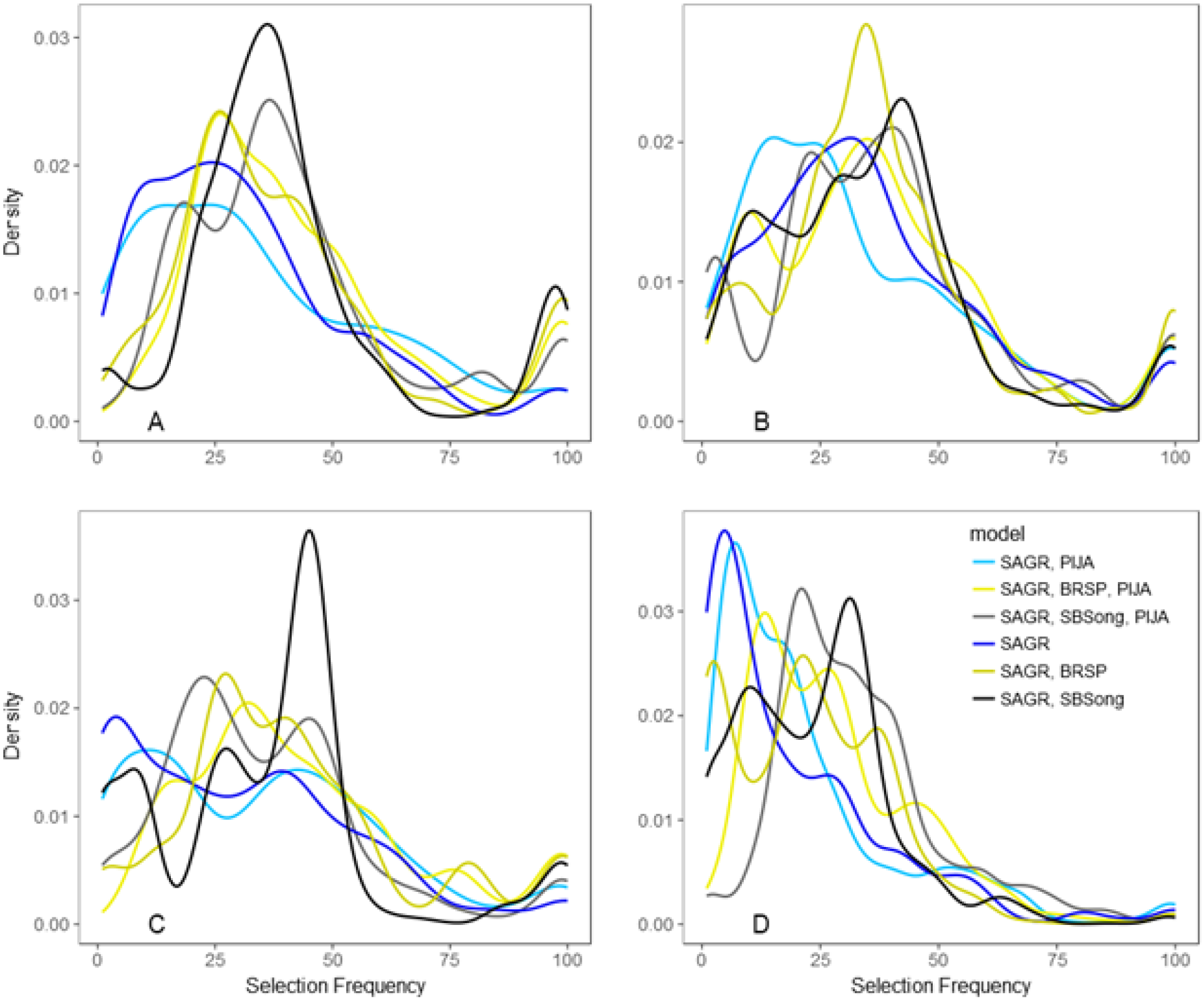
Variability in selection frequencies between Nevada (A), Oregon (B), Utah (C), and Idaho (D). Color pairings depict models with (lighter) and without pinyon jay (darker).

A comparison of areas experiencing selection frequency loss or gain as a result of the inclusion of pinyon jay to the conifer optimizations (Figure 4) with areas of predicted conifer removal occurring between 2011-2013 and 2015-2017 (Reinhardt et al. 2020) (Figure 5) indicated that a majority of estimated management efforts have occurred in areas experiencing an increase in selection frequency as a result of the inclusion of pinyon jay (sage-grouse only: 259.5 km^2^, 19.4%; sage-grouse + brewer’s sparrow: 331.5 km^2^, 24.8%; sage-grouse + all sagebrush obligates: 293.3 km^2^, 22%) or in areas experiencing little to no change in selection frequency (Δsf < ±0.5 SD) (sage-grouse only: 876.15 km^2^, 65.6%; sage-grouse + brewer’s sparrow: 828.2 km^2^, 62%; sage-grouse + all sagebrush obligates: 798.52 km^2^, 59.8%). Relatively fewer estimated conifer removal efforts occurred on areas demonstrating a loss in selection frequency as a result of the incorporation of pinyon jay (sage-grouse only: 199.9 km^2^, 14.9%; sage-grouse + brewer’s sparrow: 175.8 km^2^, 13.2%; sage-grouse + all sagebrush obligates: 243.7 km^2^, 18.2%).

**Figure 4.**
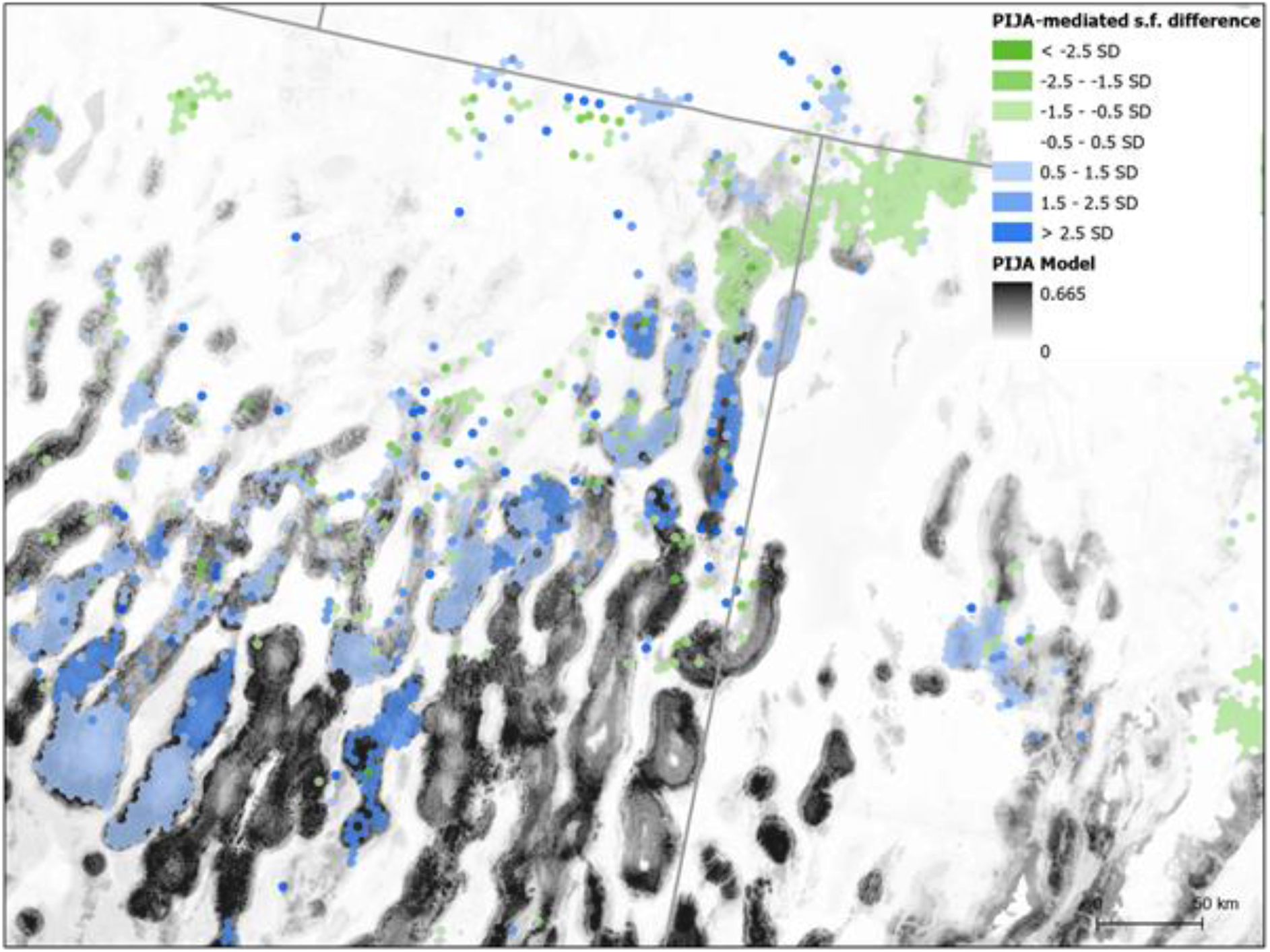
Difference in selection frequencies (s.f.) between sage-grouse + all sagebrush-obligate songbird models with without pinyon jay. Green shading depicts increasing s.f. with pinyon jays, with corresponding moderate (light) and severe (dark) decreases in blue. Grayscale represents pinyon jay habitat suitability from Tack et al. (2022).

**Figure 5.**
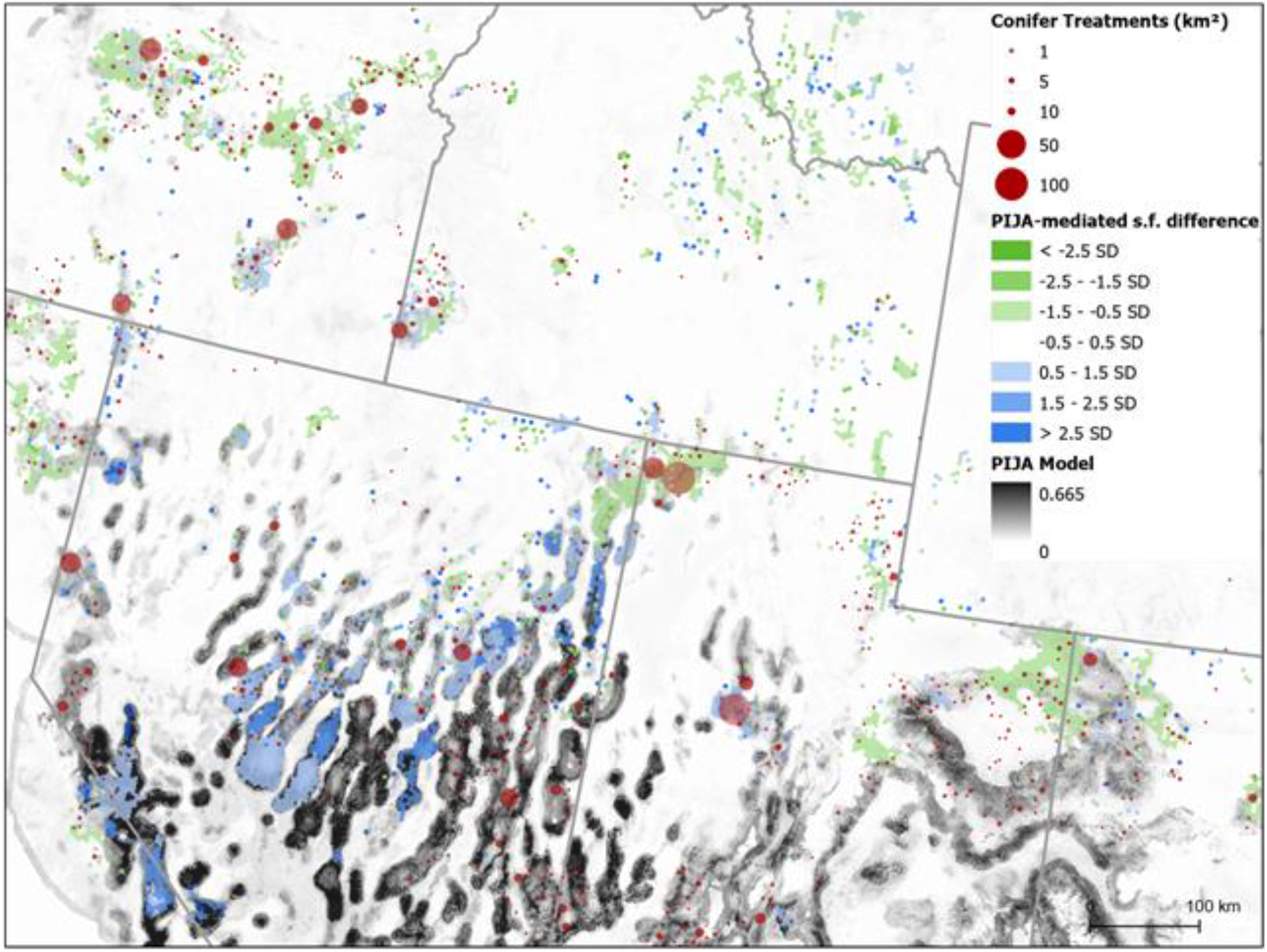
Conifer removal treatments identified by Reinhardt et al. (2020) are compared with differences in selection frequency (s.f.) between sage-grouse + all sagebrush-obligate songbird models including and excluding pinyon jay. Removals are dark red circles, with larger circles representing larger treatments. Most (81.8%) treatments are outside of highly selected pinyon jay areas (in blue), suggesting that ongoing management largely avoids pinyon jay strongholds.

## Discussion

Here, we provide a set of spatial optimizations of areas for conifer removal across the sage-grouse range which balances the habitat requirements of multiple species, some of which share habitat overlap with the sage-grouse and others which demonstrate competing habitat requirements. Total area considered in optimization output ranges from 79,072 km^2^ in the sage-grouse only scenario to over 100,000 km^2^ in the sage-grouse + all sagebrush-obligates scenario. Indeed, adding sagebrush-obligate songbirds to the optimization scenarios resulted in an across-the-board increase in areas selected by the algorithm (Table 2). While a substantial change, this is an unsurprising result because while the sagebrush obligate songbirds likely share many important habitat characteristics with sage-grouse, they are unlikely to demonstrate perfect overlap (see for e.g., Rowland et al. 2006). Adding additional sagebrush obligates to the optimization scenario was therefore likely to expand the area selected to meet the optimization parameters.

The incorporation of pinyon jay resulted in substantial changes to overall optimization results, changing the distribution of priority areas overall (Figure 2) and between states (Table 2, Figures 1 & 3). This shift in priority reflects the competing habitat requirements of pinyon jay and the sagebrush obligate species. Areas most important to pinyon jay – comprising most of the species’ range – were largely located in the Central Basin and Range Ecoregion, particularly Nevada and Utah. It was in this region where the largest apparent impacts of incorporating pinyon jay on the conifer removal optimizations was apparent (Figure 4), with several large areas demonstrating a loss in priority when pinyon jay was incorporated into the optimization scenario.

The areas where the inclusion of pinyon jay results in a meaningful loss in priority (Figure 4) do not necessarily imply conifer removal should be avoided, but instead these areas represent places where a more nuanced and holistic approach to conifer removal is likely required to balance the needs of sagebrush obligates with pinyon jay. The long-term population decline of the pinyon jay (Sauer et al. 2017; Tack et al. *in review*) coincides with the long-term expansion and infill of native conifer woodlands across the intermountain west (Miller et al. 2008, Fillippelli et al. 2020), suggesting that habitat quantity may not be a driving factor in population decline (Boone et al. 2018). Boone et al. (2018) suggest that factors such as pinyon pine growth and productivity, aspects of habitat quality, landscape structural changes, and climate change may be driving pinyon jay population declines, and some of these factors may be impacted by conifer removal efforts in the areas identified by our analyses. Given current gaps in our knowledge about how pinyon jay responds to conifer management, coupling co-produced science and monitoring (Naugle et al. 2020) with woodland restoration efforts in the Central Basin and Range Ecoregion seems prudent to addressing changing woodland conditions (Filippelli et al. 2020, Shriver et al. 2021) in an adaptive management framework.

Our comparison of the spatially explicit effects of pinyon jay on the optimization scenarios in this study with recent areas of conifer removal as estimated by Reinhardt et al. (2020) revealed that a small proportion (13-18%) of management efforts had occurred on areas we surmise as being important for pinyon jay (Figure 5). The large majority of management efforts appear to be occurring in areas we predict as being less important to pinyon jay (i.e., areas where incorporating pinyon jay results in a gain in priority, or little to no change). This result is encouraging and falls in line with other recent work demonstrating similar results (Tack et al. *in review*). Considered together, these results should indicate that the ongoing management of encroaching conifer woodlands in sagebrush ecosystems across the intermountain west is largely avoiding areas that may be particularly important for pinyon jay. This result may provide additional context for the ongoing debate regarding sage-steppe habitat improvement efforts, and the question as to whether pinyon-juniper removal efforts are resulting in widespread loss of habitat for pinyon-juniper woodland-associated species (Boone et al. 2018; Coe et al. 2018; Maestas et al. 2019; Clark et al. 2019). More broadly, our optimization results suggest an abundance of opportunities for sagebrush habitat restoration in areas outside pinyon jay strongholds (Figures 1 & 5).

Optimization results incorporating all species (Figure 1) suggest that some of the highest priority areas remain consistent whether pinyon jay is included in the assessment or not. This includes large parts of central and southeastern Oregon, the northeastern corner of California, northern Utah, and Idaho – particularly southwest Idaho. These high priority areas also happen to be some of the big ‘hotspot’ regions in terms of quantified ongoing conifer removal efforts (Figure 5), suggesting that the broad collaborative effort to restore sagebrush ecosystems and remove encroaching conifer across much of the west has so far successfully targeted some of the best-available areas for conifer removal as indicated by our optimizations.

In addition to the important implications of our analyses with regard to conifer removal priority, sagebrush obligate bird species, and pinyon jay, the optimizations created here have the potential to be useful as decision support tools when considering and planning conifer removal efforts across the western U.S. These results, while spatially explicit, are intentionally coarse (16.24 km^2^ hexagonal grid) because they seek to analyze large-scale patterns in conifer optimization across the intermountain west while incorporating multiple species with differing habitat requirements. Local-level assessments of ecological site potential and conditions are still required to determine appropriateness of conifer removal treatments and risks (e.g., invasive annual grasses). The systematic conservation planning approach that the optimizations in this study employed, however, are quite adaptable; this approach could be extended or modified to include additional bird species or other taxa or to consider management goals beyond conifer removal. Indeed, we hope that the approach outlined here will provide a useful blueprint for future analyses.

This work builds on previous and ongoing research concerned with sage-grouse conservation, sagebrush ecosystem restoration, and a growing focus on restoration that can realize multi-species benefits. The optimizations conducted here seek to evaluate the potential differences in conifer removal priority for specific areas as a result of the inclusion of species beyond the sage-grouse – including other sagebrush obligates as well as a species with competing habitat requirements in the pinyon jay. Consideration of taxa beyond birds is likely to be important going forward, as sagebrush ecosystem management efforts become increasingly community focused. The continued integration of new spatial datasets into management optimizations will arm managers on the ground with the tools necessary to ensure conservation goals are met for co-occurring species with divergent habitat requirements.

## Acknowledgements

The Western Association of Fish and Wildlife Agencies provided support through the Sagebrush Science Initiative. The findings and conclusions in this article are those of the authors and should not be construed to represent any official USDA or U.S. Government determination or policy.

## Role of the funding source

This study was funded by the Sagebrush Science Initiative, a collaboration between the Western Association of Fish and Wildlife Agencies and US Fish and Wildlife Service, the US Bureau of Land Management, and the US Department of Agriculture’s Natural Resource Conservation Service, Sage Grouse Initiative. Funding sources were not involved in the data collection, study design, analyses, or interpretation.

## Notes

### Competing Interest Statement

The authors have declared no competing interest.

## References

Allred, B. W., B. T. Bestelmeyer, C. S. Boyd, C. Brown, K. W. Davies, M. C. Duniway, L. M. Ellsworth, T. A. Erickson, S. D. Fuhlendorf, T. V. Griffiths, V. Jansen, M. O. Jones, J. Karl, A. Knight, J. D. Maestas, J. J. Maynard, S. E. McCord, D. E. Naugle, H. D. Starns, D. Twidwell, and D. R. Uden. 2021. Improving Landsat predictions of rangeland fractional cover with multitask learning and uncertainty. Methods in Ecology and Evolution 12:841–849.

Ardron, J. A., Possingham, H. P and C. J. Klein (Editors). 2010. Marxan good practices handbook. Version 2. Pacific Marine Analysis and Research Association, Victoria, British Columbia, Canada.

Baruch-Mordo, S., J. S. Evans, J. P. Severson, D. E. Naugle, J. D. Maestas, J. M. Kiesecker, M. J. Falkowski, C. A. Hagen, and K. P. Reese. 2013. Saving sage-grouse from the trees: A proactive solution to reducing a key threat to a candidate species. Biological Conservation 167: 233–241.

Bates, J. D. 2005. Herbaceous response to cattle grazing following juniper cutting in Oregon. Rangeland Ecology and Management 58: 225–233.

Bates, J. D., K. W. Davies, A. Hulet, R. F. Miller, and B. Roundy. 2017. Sage-grouse groceries: Forb response to piñ on-juniper treatments. Rangeland Ecology and Management 70:106–115.

Bender, L. C., L. A. Lomas, and T. Kamienski. 2007. Habitat effects on condition of doe mule deer in arid mixed woodland-grassland. Rangeland Ecology and Management 60:277–284.

Bergman, E. J., C. J. Bishop, D. J. Freddy, G. C. White, and P. F. Doherty, Jr. 2014a. Habitat management influences overwinter survival of mule deer fawns in Colorado. Journal of Wildlife Management 78:448–455.

Bergman, E. J., P. F. Doherty Jr., C. J. Bishop, L. L. Wolfe, and B. A. Banulis. 2014b. Herbivore body condition response in altered environments: mule deer and habitat management. PLoS One 9:e106374.

Beyer, H. L., Y. Dujardin, M. E. Watts, and H. P. Possingham. 2016. Solving conservation planning problems with integer linear programming. Ecological Modelling 328:14–22.

Bombaci, S. and L. Pejchar. 2016. Consequences of pinyon and juniper woodland reduction for wildlife in North America. Forest Ecology and Management 365:34–50.

Boone, J. D., E. Ammon, and K. Johnson, 2018. Long-term declines in the Pinyon Jay and management implications for piñ on–juniper woodlands, in Trends and traditions: Avifaunal change in western North America (W. D. Shuford, R. E. Gill Jr., and C. M. Handel, editors.). Pages 190–197 In Studies of Western Birds 3. Western Field Ornithologists, Camarillo, California.

Carwardine, J. W. A. Rochester, K. S. Richardson, K. J. Williams, R. L. Pressey, and H. P. Possingham. 2007. Conservation planning with irreplaceability: Does the method matter? Biodiversity and Conservation 16:245–258.

Chambers, J. C., B. A. Roundy, R. R. Blank, S. E. Meyer, and A. Whittaker. 2007. What makes Great Basin sagebrush ecosystems invasible by Bromus tectorum? Ecological Monographs 77:117–145.

Chambers, J. C., Maestas, J. D., Pyke, D. A., Boyd, C. S., Pellant, M., and Wuenschel, A., 2017. Using resilience and resistance concepts to manage persistent threats to sagebrush ecosystems and greater sage-grouse. Rangeland Ecology & Management, 70(2), 149–164.

Chambers, J. C., Miller, R. F., Board, D. I., Pyke, D. A., Roundy, B. A., Grace, J. B., … and Tausch, R. J., 2014. Resilience and resistance of sagebrush ecosystems: implications for state and transition models and management treatments. Rangeland Ecology & Management, 67(5), 440–454.

Chambers, J. C., R. F. Miller, D. I. Board, D. A. Pyke, B. A. Roundy, J. B. Grace, E. W. Schupp, and R. J. Tausch. 2014. Resilience and resistance of sagebrush ecosystems: Implications for state and transition models and management treatments. Rangeland Ecology and Management 67:440–454.

Chambers, J.C.; J. D. Maestas, D. A. Pyke, C. Boyd, M. Pellant, and A. Wuenschel. 2017. Using resilience and resistance concepts to manage persistent threats to sagebrush ecosystems and greater sage-grouse. Rangeland Ecology and Management 70:149–164.

Clark, D. A., Coe, P. K., Gregory, S. C., Hedrick, M. J., Johnson, B. K., and D. H. Jackson. 2019. Habitat use informs species needs and management: A Reply to Maestas et al. Journal of Wildlife Management, 83:762–766.

Coe, P. K., D. A. Clark, R. M. Nielson, S. C. Gregory, J. B. Cupples, M. J. Hedrick, B. K. Johnson, and D. H. Jackson. 2018. Multiscale models of habitat use by mule deer in winter. Journal of Wildlife Management 82:1285–1299.

Crow, C. and C. Van Riper. 2010. Avian community responses to mechanical thinning of a pinyon-juniper woodland: Specialist sensitivity to tree reduction. Natural Areas Journal 30:191–201.

Daly, C., M. Halbleib, J. I. Smith, W. P. Gibson, M. K. Doggett, G. H. Taylor, J. Curtis, and P. P. Pasteris. 2008. Physiographically sensitive mapping of climatological temperature and precipitation across the conterminous United States. International Journal of Climatology.. 28:2031–2064.

Doherty, K. E., J. S. Evans, P. S. Coates, L. Juliusson, and B. C. Fedy. 2016. Importance of regional variation in conservation planning: a rangewide example of the greater sage-grouse. Ecosphere 7:e01462.

Donnelly, J. P., J. D. Tack, K. E. Doherty, D. E. Naugle, B. W. Allred, and V. J. Dreitz. 2017. Extending conifer removal and land protection strategies from sage-grouse to songbirds, a range-wide assessment. Rangeland Ecology and Management 70:95–105.

Falkowski, M. J., J. S. Evans, D. E. Naugle, C. A. Hagen, S. A. Carleton, J. D. Maestas, A. H. Khalyani, A. J. Poznanovic, and A. J. Lawrence. 2017. Mapping tree canopy cover in support of proactive prairie grouse conservation in western North America. Rangeland Ecology and Management 70:15–24.

Falkowski, M. J., Smith, A. M. S., Hudak, A. T., Gessler, P. E., Vierling, L. A. and N. L. Crookston. 2006. Automated estimation of individual conifer tree height and crown diameter via two-dimensional spatial wavelet analysis of lidar data. Canadian Journal of Remote Sensing 32:153–161.

Fick, S. E., T. W. Nauman, C. C. Brungard, and M. C. Duniway. 2022. What determines the effectiveness of Pinyon-Juniper clearing treatments? Evidence from the remote sensing archive and counter-factual scenarios. Forest Ecology and Management 505:119879.

Filippelli, S.K., Falkowski, M.J., Hudak, A.T., Fekety, P.A., Vogeler, J.C., Khalyani, A.H., Rau, B.M. and Strand, E.K., 2020. Monitoring pinyon-juniper cover and aboveground biomass across the Great Basin. Environmental Research Letters, 15(2), p.025004.

Gurobi Optimization I. 2017. Gurobi optimizer reference manual. Retrieved from http://www.gurobi.com.

Hanson, J. O., R. Schuster, N. Morrell, M. Strimas-Mackey, M.E. Watts, P. Arcese, J. Bennett, and H. P. Possingham. 2019. prioritizr: Systematic Conservation optimization in R. R package version 4.0.2.14. Available at https://github.com/prioritizr/prioritizr.

Holmes, A. L., J. D. Maestas, and D. E. Naugle. 2017. Non-game bird responses to removal of western juniper in sagebrush-steppe. Rangeland Ecology and Management 70:87–94.

Homer, C. G., J. Dewitz, L. Yang, S. Jin, P. Danielson, C. J. Xian, N. Herold, J. Wickham, and K. Megown. 2015. Completion of the 2011 National Land Cover Database for the conterminous United States – representing a decade of land cover change information, Photogrammetric Engineering and Remote Sensing 81: 345–353.

Jones M. O., B. W. Allred, D. E. Naugle, J. D. Maestas, P. Donnelly, L. J. Metz, J. Karl, R. Smith, B. Bestelmeyer, C. Boyd, J. D. Kerby, and J.D. McIver. 2018. Innovation in rangeland monitoring: annual, 30 m, plant functional type percent cover maps for U.S. rangelands, 1984-2017. Ecosphere 9:e02430.

Kottek, M., J. Grieser, C. Beck, B. Rudolf, and F. Rubel. 2006. World map of the Köppen-Geiger climate classification updated. Meteorologische Zeitschrift 15:259–263.

Maestas, J. D., C. A. Hagen, J. T. Smith, J. D. Tack, B. Allred, T. Griffiths, C. Bishop, and D. E. Naugle. 2019. Mule deer juniper use is an unreliable indicator of habitat quality: Comments on Coe et al. (2018). The Journal of Wildlife Management 83:755–762.

Maestas, J. D., D. E. Naugle, J. C. Chambers, J. D. Tack, C. S. Boyd, and J. M. Tague, 2021, Chapter M. Conifer Expansion, in T. E. Remington, P. A. Deibert, S. E. Hanser, D. M. Davis, L. A. Robb, and J. L. Welty, eds., Sagebrush conservation strategy—Challenges to sagebrush conservation: U.S. Geological Survey Open-File Report 2020–1125: doi:https://doi.org/10.3133/ofr20201125.

Miller, R. F., D. E. Naugle, J. D. Maestas, C. A. Hagen, and G. Hall (2017). Special Issue: Targeted Woodland Removal to Recover at-Risk Grouse and Their Sagebrush-Steppe and Prairie Ecosystems. Rangeland Ecology & Management 70:1–8.

Miller, R. F., J. C. Chambers, and M. Pellant. 2014. A field guide to selecting the most appropriate treatments in sagebrush and pinyon-juniper ecosystems in the Great Basin: evaluating resilience to disturbance and resistance to invasive annual grasses and predicting vegetation response. Fort Collins, CO, USA: U.S. Department of Agriculture, Forest Service, RMRS-GTR-322.

Miller, R. F., J. C. Chambers, L. Evers, C. J. Williams, K. A. Snyder, B. A. Roundy, and F. B. Pierson. 2019. The ecology, history, ecohydrology, and management of pinyon and juniper woodlands in the Great Basin and Northern Colorado Plateau of the western United States. General Technical Report RMRS-GTR-403, Rocky Mountain Research Station, Fort Collins, Colorado, United States Forest Service.

Miller, R. F., R. J. Tausch, E. D. McArthur, D. D. Johnson, and S. C. Sanderson. 2008. Age structure and expansion of pinyon-juniper woodlands: A regional perspective in the Intermountain West. U.S. Department of Agriculture, Forest Service, Rocky Mountain Research Station, Fort Collins, Colorado.

Miller, R. F., S. T. Knick, D. A. Pyke, C. W. Meinke, S. E. Hanser, M. J. Wisdom, and A. L. Hild. 2011. Characteristics of sagebrush habitats and limitations to long-term conservation. Pages 145–184 in S. T. Knick and J. W. Connelly, editors. Greater sage-grouse: ecology and conservation of a landscape species and its habitats. Studies in Avian Biology 38. University of California Press, Berkeley, California.

Miller, R. F., T. J. Svejcar, and J. A. Rose. 2000. Impacts of western juniper on plant community composition and structure. Journal of Range Management 53:574–585.

Nackley, L. L., A. G. West, A. L. Skowno, and W. J. Bond. 2017. The nebulous ecology of native invasions. Trends in Ecology and Evolution 32: 814–824.

Naugle, D. E., Allred, B. W., Jones, M. O., Twidwell, D., and Maestas, J. D., 2020. Coproducing science to inform working lands: The next frontier in nature conservation. BioScience, 70(1), 90–96.

Olsen, A. C., J. P. Severson, J. D. Maestas, D. E. Naugle, J. T. Smith, J.D. Tack, K. H. Yates, and C. A. Hagen. 2021. Reversing tree expansion in sagebrush steppe yields population-level benefit for imperiled grouse. Ecosphere 12:e03551.

Prochazka, B. G., P. S. Coates, M. A. Ricca, M. L. Casazza, K. B. Gustafson, and J. M. Hull. 2017. Encounters with pinyon-juniper influence riskier movements in greater sage-grouse across the Great Basin. Rangeland Ecology and Management 70:39–49.

R Core Team. 2019. R: A language and environment for statistical computing. R Foundation for Statistical Computing, Vienna, Austria. Available at http://www.R-project.org

Rabon, J. C., P. S. Coates, M. A. Ricca, and T. N. Johnson. 2021. Does reproductive status influence habitat selection by female greater sage-grouse in a sagebrush-juniper landscape? Rangeland Ecology and Management 79:150–163.

Reinhardt, J. R., D. E. Naugle, J. D. Maestas, B. Allred, J. Evans, and M. Falkowski. 2017. Next-generation restoration for sage-grouse: a framework for visualizing local conifer cuts within a landscape context. Ecosphere 8:e01888.

Reinhardt, J. R., S. Filippelli, M. Falkowski, B. Allred, J. D. Maestas, J. C. Carlson, and D. E. Naugle. 2020. Quantifying pinyon-juniper reduction within North America’s sagebrush ecosystem. Rangeland Ecology and Management 73:420–432.

Rickart, E.A., Robson, S.L., Heaney, L.R., 2008. Mammals of Great Basin National Park, Nevada: Comparative field surveys and assessment of faunal change. Western North American Naturalist 4:77–114.

Roundy, B. A., R. F. Miller, R. J. Tausch, K. Young, A. Hulet, B. Rau, B. Jessop, J. C. Chambers, and D. Eggett. 2014. Understory cover responses to piñ on-juniper treatments cross tree dominance gradients in the Great Basin. Rangeland Ecology and Management 67:482–494.

Rowland, M., Wisdom, M., Suring, L. and C. Meinke. 2006. Greater sage-grouse as an umbrella species for sagebrush-associated vertebrates. Biological Conservation 129:323–335.

Sauer, J. R., Niven, D. K., Hines, J. E., Ziolkowski, D. J., Jr., Pardieck, K. L., Fallon, J. E., and Link, W. A. 2017. The North American Breeding Bird Survey, results and analysis 1966–2015, version 02.07.2017. USGS Patuxent Wildlife Research Center, Laurel, MD; http://www.mbr-pwrc.usgs.gov/bbs/.

Severson, J. P. 2016. Greater sage-grouse response to conifer encroachment and removal. Dissertation, University of Idaho, Moscow, Idaho.

Severson, J. P., C. A. Hagen, J. D. Maestas, D. E. Naugle, J. T. Forbes, and K. P. Reese. 2017a. Short-term response of sage-grouse nesting to conifer removal in the northern Great Basin. Rangeland Ecology & Management 70:50–58.

Severson, J. P., C. A. Hagen, J. D. Tack, J. D. Maestas, D. E. Naugle, J. T. Forbes, and K. P. Reese. 2017b. Better living through conifer removal: a demographic analysis of sage-grouse vital rates. PLoS One 12:e0174347.

Shriver, R. K., C. B. Yackulic, D. M. Bell, and J. B. Bradford. 2021. Quantifying the demographic vulnerabilities of dry woodlands to climate and competition using rangewide monitoring data. Ecology 102(8):e03425. 10.1002/ecy.3425

Tack, J., Smith, J.T., Doherty, K.E., Donnelly, P.J., Maestas, J.D., Allred, B.W., Reinhardt, J.R., Morford, S.L. and Naugle, D.E., in review. Regional context for balancing sagebrush- and woodland-dependent songbird needs with targeted pinyon-juniper management.

United States Fish and Wildlife Service (USFWS). 2013. Greater sage-grouse (Centrocercus urophasianus) conservation objectives Final Report. United States Department of the Interior, Denver, Colorado.

United States Fish and Wildlife Service (USFWS). 2015. Endangered and threatened wildlife and plants; 12-month finding on a petition to list greater sage-grouse (Centrocercus urophasianus) as an endangered or threatened species. Federal Register 80:59858–59942.

Zeller, K.A., Cushman, S.A., Van Lanen, N.J., Boone, J.D. and Ammon, E., 2021. Targeting conifer removal to create an even playing field for birds in the Great Basin. Biological Conservation, 257. https://doi.org/10.1016/j.biocon.2021.109130

